# High-gamma activity in the human hippocampus during inter-trial rest periods of a virtual navigation task

**DOI:** 10.1101/252288

**Authors:** Yi Pu, Brian R. Cornwell, Douglas Cheyne, Blake W. Johnson

## Abstract

In rodents, hippocampal cell assemblies formed during learning of a navigation task are observed to re-emerge during resting (offline) periods, accompanied by high-frequency oscillations (HFOs). This phenomenon is believed to reflect mechanisms for strengthening newly-formed memory traces. Using magnetoencephalography recordings and a beamforming source location algorithm (synthetic aperture magnetometry), we investigated high-gamma (80 – 140 Hz) oscillations in the hippocampal region in 18 human participants during inter-trial rest periods in a virtual navigation task. We found right hippocampal gamma oscillations mirrored the pattern of theta power in the same region during navigation, varying as a function of environmental novelty. Gamma power during inter-trial rest periods was positively correlated with theta power during navigation in the first training set when the environment was new and predicted faster learning in the subsequent training set two where the environment became familiar. These findings provide evidence for human hippocampal reactivation accompanied by high-gamma activities immediately after learning and establish a link between hippocampal high-gamma activities and memory consolidation.

## Introduction

The formation of spatial memories is proposed to proceed in two stages (Buzsaki, 1989, 2015). In the encoding phase, during active exploration of an environment, a transient change of synaptic strengths in the hippocampus is formed accompanied by theta-band neuronal oscillations. Subsequently, during ‘offline’ states, including slow-wave sleep and quiet wakefulness, the newly formed synaptic network re-emerges, accompanied by high frequency oscillations (HFOs), operating to potentiate and strengthen the synaptic changes and thereby consolidate the otherwise labile memory traces.

Rodent studies have shown that the sequential activation of place cells during navigation reoccurs (“replays”) when the animal is asleep or in a state of awake immobility after exploration, and this replay is accompanied by HFOs (O’Neill et al., 2010). Disruption of hippocampal HFOs impairs spatial learning (Ego-Stengel & Wilson, 2010; Girardeau et al., 2009; Jadhav et al., 2012), suggesting a causal relationship between HFOs and memory formation. Replay is sensitive to environmental novelty (Carr et al., 2011): After navigating in a new environment, the strength of place cell replay is stronger (Diba & Buzsaki, 2007; O’Neill et al., 2008) and the probability of the occurrence of HFOs and the firing rates of place cells are significantly higher (Cheng & Frank, 2008) than that following navigation in a familiar environment.

The two-stage model has been intensively investigated in animal models. Are comparable neurophysiological learning mechanisms used in the human hippocampus? Currently, there is very limited, but highly suggestive evidence that this is the case. fMRI studies (Deuker et al., 2013; Gruber et al., 2016; Staresina et al., 2013; Tambini & Davachi, 2013; Tambini et al., 2010) have reported that brain regions that are active during learning, are reactivated during sleep or rest periods after learning. For instance, using multivariate pattern classification analysis, Deuker et al. (2013) found stimulus-specific patterns during encoding reoccurred spontaneously during postlearning resting periods and sleep.

To date, there was only limited electrophysiological evidence pertaining to the two-stage model. For instance, from intracranial recordings in human patients, Axmacher et al. (2008) reported robust high-gamma rhythms (80 – 140 Hz) during the post-learning sleep period in the hippocampus and rhinal cortex, and high-gamma in the rhinal cortex was positively correlated with subsequent memory performance. Using noninvasive MEG measurements, Cornwell et al., (2014) reported that post-learning high-gamma power was positively correlated with spatial learning performance before the rest period. But in the two studies, there was no control condition, it is therefore uncertain whether the high-gamma activities reported are learning-specific or only a general trait marker related to general cognitive processing speed. Recently, using decoding methods, Kurth-Nelson et al. (2016) found during object-free periods after learning, the brain spontaneously replayed the representations of four objects learned in the learning period in a reverse order. This study reveals that the replay mechanism might be a fundamental neural computation in human brain as well. However, these results are on the MEG sensor level and which source brain regions and neuronal oscillations are related to the replay phenomenon is unknown.

In the present study, we leveraged the high time resolution of MEG to investigate the temporal dynamics of human hippocampal “reactivation” during ITI immediately after learning trials. MEG was recorded while participants performed two training sets of a virtual Morris water maze task. Each set included a *hidden platform* condition (task: finding the hidden platform) and a *random swimming* condition (task: aimlessly swimming in a pool without platforms). Environment layouts of each condition in the two training sets were the same. In a previous report on data from the same experiment (Pu et al., 2017), we studied the role of low frequency theta oscillations (4 – 8 Hz) in spatial encoding during navigation. We found that there was significantly greater theta power in right hippocampus in the first compared to the second training set, which was associated with environment encoding; and there was significantly more left hippocampal theta in the *hidden platform* condition than in the *random swimming* condition, which was associated with encoding of the hidden platform location.

Here we asked whether hippocampal high-gamma power during ITI would reflect the patterning of hippocampal theta power change during navigation, and whether the power of high-gamma was correlated with the theta power, since replay is proportional to previous learning in rodents (Sutherland & McNaughton, 2000, see Buzsaki, 2015 for a review). We also investigated whether high-gamma power after navigating in the new environment (first training set) was associated with speed of spatial learning in the familiar environment (second training set), since consolidation of newly-learned environment to form a cognitive map of the space should facilitate flexible navigation to new locations in the same environment (Wolbers & Hegarty, 2010).

## Materials and Methods

### Participants

Eighteen male participants (mean age = 29 years; range = 18 – 39 years) participated in the study. Two additional participants were excluded from the final data analyses because of the excessive head movement. The study was approved by Macquarie University’s human subjects ethics committee. All participants gave written informed consent. Analysis of data during active navigation was previously reported in Pu et al. (2017). The current analysis investigated high-gamma during the ITI of the experiment when participants rested quietly following each trial of spatial navigation.

### Experiment design

A detailed description of the experimental paradigm is in Pu et al. (2017). In brief, naive participants performed two training sets of a virtual Morris water maze task. In each training set of the task, there were two conditions. In the *hidden platform* condition, participants needed to find a hidden platform submerged in opaque water by using the visual cues on the walls surrounding the virtual pool. In the *random swimming* condition, participants moved aimlessly in the same virtual pool (but with no visual cues on the walls). The environment of each condition in the two training sets was the same, thus the environment in the first training set was defined as new environment and that in the second one as familiar environment. Therefore, the difference between the two training sets allowed us to measure *learning of the environment* (Pu et al., 2017), and the difference between *hidden platform* condition and *random swimming* condition provided an index of *goal-directed spatial navigation* (Cornwell et al., 2008). To avoid the possibility that environment learning was confounded with learning a specific location, the location of the hidden platform was changed and counterbalanced between the training sets.

In each training set, there were 40 trials including 20 hidden platform and 20 random swimming trials respectively, presented in alternating blocks of four trials. Between each trial, there was a 4.5 – 5.5 s ITI (Figure 1), during which a gray screen was presented and participants rested quietly without movement.

**Figure 1.**
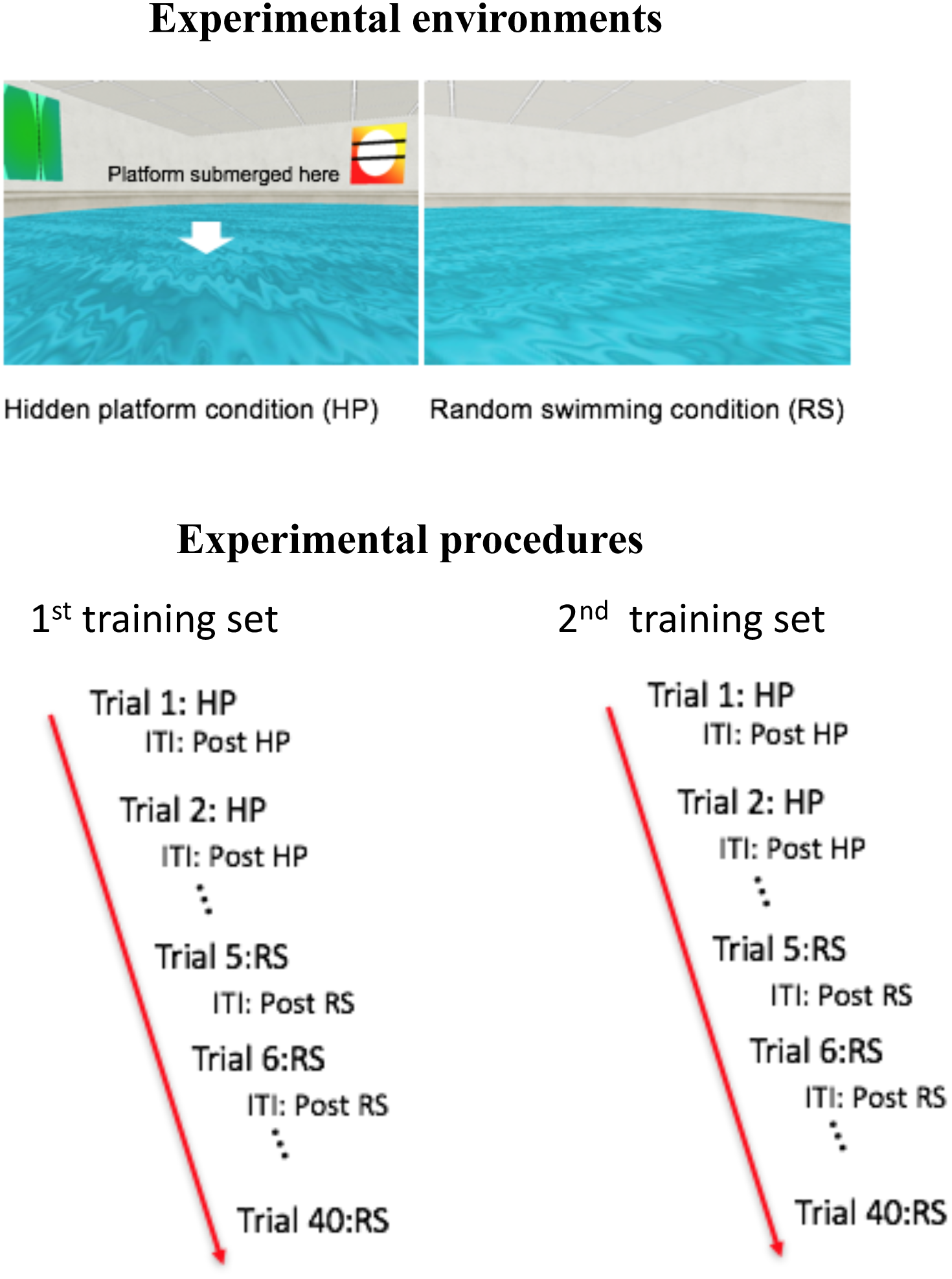
Overview of the experiment. The upper panel shows environments of the two conditions. The lower panel shows a flow chart of experimental procedure. In each training set, there were 40 trials, including 20 hidden platform trials (the task was to find the hidden platform in a pool with four cues) and 20 random swimming trials (the task was to aimless swimming in a pool without visual cues and platform), which were alternatively presented (4 hidden platform trials, 4 random swimming trials, 4 hidden platform trials, 4 random swimming trials...). The interval between each trial (ITI) was 4.5 – 5.5s, during which a grey screen was presented and participants rested quietly without movement. HP: hidden platform condition. RS: random swimming condition.

### Behavioral measures

The length of the path taken from the starting position to the hidden platform in each trial was recorded. Learning rate was computed as the average path length of the first block minus that of the last one, divided by the number of blocks. To capture a more instant learning change, the slope of a linear regression line fit to the average path lengths across the five blocks was computed.

### MEG recordings

Recordings were made in a magnetically shielded room (Fujihara Co. Ltd., Tokyo, Japan) with a 160-channel KIT system (Model PQ1160R-N2, Kanazawa, Japan) with superconducting quantum interference device (SQUID)-based first-order axial gradiometers (50-mm baseline; Kado et al., 1999; Uehara et al., 2003). Neuromagnetic signals were digitized continuously at a sampling rate of 1000 Hz filtered at 0.03 and 200 Hz. Before recordings, the locations of the five marker coils and three fiducial markers, and the participant’s head shape were digitised with a pen digitizer (Polhemus Fastrack, Colchester, VT, USA). The five marker coils were energized before and after each training set to determine head movement and position within the MEG dewar.

### MRI scans

High-resolution T1-weighted anatomical magnetic resonance images (MRIs) were acquired in a separate session at Macquarie University Hospital, using a 3T Siemens Magnetom Verio scanner with a 12-channel head coil. Images were obtained using 3D GR\IR scanning sequence with the following parameters: repetition time, 2000 ms; echo time, 3.94 ms; flip angle, 9 degrees; slice thickness, 0.93 mm; field of view, 240 mm; image dimensions, 512 × 512 × 208.

## MEG analyses

### High-gamma during ITI

The MEG data during the ITI were epoched (-4.5 – 0 s; 0 s was the onset of the next trial; 4.5 s was the shortest ITI across trials) and were labeled as *post hidden platform* condition and *post random swimming* condition respectively. Sources were reconstructed using synthetic aperture magnetometry (SAM) beamformer analysis (Hillebrand et al., 2005; Robinson & Vrba, 1999) implemented in the BrainWave toolbox (version 3.0, http://cheynelab.utoronto.ca). SAM was performed on unaveraged data so that it can identify sources that are not phase-locked or time-locked activities. It estimates power changes within specific frequency ranges and time windows across the whole brain without a prior assumption of the number and the locations of the active source (Robinson & Vrba, 1999).

MEG has been shown to reliably localize activity from the hippocampus in both simulation studies (e.g., Attal et al., 2007; Chupin et al., 2002; Meyer et al., 2017; Quraan et al., 2011; Stephen et al., 2005) and empirical experiments (e.g., Backus et al., 2016; Cornwell et al., 2008; Riggs et al., 2009; Tesche & Karhu, 2000). Recently, Crespo-Garcia et al. (2016) have shown an agreement between simultaneous intracranial depth recordings and MEG virtual sensor recordings of hippocampal activity.

Due to the 1/f power function of the MEG, high-gamma signal power is in general much smaller than power at lower frequencies. Nevertheless, many previous studies have shown that high-gamma activity in many brain regions is successfully detected by MEG (e.g., Cheyne et al., 2008; Cheyne & Ferrari, 2013; Cornwell et al., 2014; Muthukumaraswamy, 2013 for a review). The hippocampus in particular is known to be a robust generator of gamma oscillations: Invasive recordings in animal models have shown substantially *greater* power of high frequency gamma during rest/sleep compared to low frequency theta during navigation (Buzsaki, 2015; Buzsaki & Silva, 2012).

Beamformer source reconstruction is achieved by first defining a source space of volumetric grids encompassing the whole head. SAM operates by constructing an adaptive spatial filter (beamformer weights) for each grid location, based on a combination of lead fields calculated from the forward solution and the data covariance matrix. Beamformer weights are convolved with the MEG sensor data to obtain a source signal for each grid element. Since the output of the spatial filter contains both the signal of interest and noise, it is necessary to estimate the noise level and normalize the output beamformer signal to obtain a relatively ‘pure’ neural signal. One commonly used method of normalization (e.g. Cornwell et al., 2014; Perry, 2015). uses a pseudo-Z metric, which divides the absolute source power of a single state by a noise estimate (Robinson & Vrba, 1999; Vrba & Robinson, 2001). Another normalization approach (e.g. Cornwell et al., 2012; Isabella et al., 2015) uses a pseudo-F or pseudo-T metric, which computes the percentage change (pseudo-F) or absolute change (pseudo-T) of the signal power in an active state relative to a control state so as to implicitly control the noise level (under the assumption that the two states have similar noise levels).

In the present analyses, the source power of high-gamma activity during the ITI rest period was computed using a pseudo-Z metric because we were interested in estimating spontaneous high-gamma power (as opposed to event-related power changes). The forward model was a single sphere volume conduction model (Lalancette et al., 2011; Sarvas, 1987) derived from individual MRIs. Data covariance matrices were calculated for the whole epoch for the frequency band of 80 – 140 Hz, the same frequency range as used in Axmacher et al. (2008) and Cornwell et al. (2014). Thus the length of the covariance matrix in the *post hidden platform* condition was 20 trials × 4.5s/trial = 90s and that in the *post random swimming* condition was 19 trials × 4.5s/trial = 85.5s. In the latter, there were 19 trials instead of 20, because the last trial of the experiment was always a random swimming trial and the experimental program aborted after the completion of the last trial. The slight difference in covariance window length for *post hidden platform* condition and *post random swimming* condition was not expected to significantly influence source estimation: Brookes et al. (2008) demonstrated that if the bandwidths of the estimated frequency band was > 50 Hz, and when covariance window length amounted to 40 s, the accuracy of source estimation would be very high and increasing the covariance window length would not greatly improve the accuracy of source estimation. Source power was estimated across the entire 3D source space at a resolution of 4×4×4 mm.

### Group statistics

The resulting volumetric SAM images were warped to a standard Talairach template space and analyzed with Analysis of Functional Neuroimaging (AFNI) software (Cox, 1996; http://afni.nimh.nih.gov/afni). To address the first research question, i.e., whether hippocampal high-gamma during ITI mirrored the pattern of hippocampal theta activity during navigation, first, we defined a region of interest (ROI) in the right hippocampus/parahippocampus, in which a significant main effect of training set was shown for theta power during navigation; and two ROIs (because this effect occurred in two time windows: 1 – 2 s and 1.5 – 2.5 s) in the left hippocampus and parahippocampus, which showed a significant main effect of condition for theta power during navigation in theta frequency band reported in Pu et al. (2017). The mean of high-gamma power (pseudo-Z values) from the above ROIs was extracted from each condition and training set. 2 (condition: post hidden platform vs. post random swimming) × 2 (training set: 1^st^ vs. 2^nd^) within-subject ANOVAs were computed for the high-gamma power in the right and left hippocampi, with the significance level corrected to p = 0.05/3 = 0.017 (because we did ANOVA analyses three times).

To examine the focality of hippocampal activations and to address the possibility that the effects seen in the ROIs were due to signal leakage from cortical regions, we performed a 2 (condition: post hidden platform vs. post random swimming) × 2 (training set: 1^st^ vs. 2^nd^) within-subject ANOVA for each voxel across the whole brain. False positives were controlled by using a small volume FDR correction method in a mask containing bilateral hippocampi and parahippocampi with the threshold of p < 0.005, q < 0.05.

### High-gamma during navigation

We also investigated whether comparable high-gamma power was elicited during the navigation period. MEG data were epoched into 0 – 4 s (0 s was the trial onset, 4 s was the fastest time from the starting point to the hidden platform among all trials and participants) for each condition (*hidden platform* and *random swimming* condition). Beamformer images were computed for the frequency range of 80 – 140 Hz for this period. Then, the standardized beamformer images were analysed with a 2 (condition: *hidden platform* vs. *random swimming*) × 2 (training set: 1st vs. 2nd) within-subject ANOVA, with the significance threshold being p < 0.005, FDR corrected q < 0.05.

### Time frequency plots

To exhibit the evolution of high-gamma power change during ITI, time frequency representations (TFRs) were constructed for the peak voxel of the hippocampal region from the whole brain analyses. A five-cycle wavelet was convolved with the beamformed source activity over a frequency range of 1 – 200 Hz in 1 Hz steps from -4.5 – 0 s using the formula of

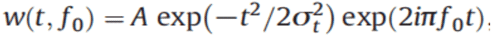

The final TFRs were presented as the power change in one condition/training set relative to the other condition/training set.

## Correlational analyses

### High-gamma during ITI versus subsequent navigation performance

To test the hypothesis that stronger replay after spatial learning in a new environment for memory consolidation of the environment is associated with quicker subsequent learning in the same environment when it became more familiar, the hippocampal high-gamma power which showed a significant effect in the ANOVA analyses in the *post hidden platform* condition relative to navigation period in *hidden platform* condition in the first training set (new environment), was correlated with the learning rate in the second training set using Pearson correlation implemented in IBM SPSS (version 23).

### High-gamma during ITI versus theta during navigation

To test the hypothesis that replay was proportional to encoding, the mean of the high-gamma power (pseudo-Z values) in the ITI after *hidden platform* condition and after *random swimming* condition in the hippocampal ROI, which showed a significant effect in the ANOVA analyses, were extracted and correlated with the mean of theta power (pseudo-Z values) in the same ROI in the time window of 1.25 – 2.25s showing a significant encoding effect in Pu et al. (2017) during navigation in the *hidden platform* condition and *random swimming* condition respectively. The method of computing pseudo-Z images for theta power during navigation was the same as used for computing pseudo-Z images for high-gamma power during ITI. To determine if the correlation was constrained within the hippocampus and parahippocampus, the mean of high-gamma power (pseudo-Z values) in the ROI was also correlated voxel-wisely with pseudo-Z image of theta power during navigation across the whole brain. The significance threshold was set as p < 0.005 (uncorrected).

### Post-hoc analyses

Since the ITI was random jittered, and in the primary analyses we epoched the data from -4.5 – 0s, to confirm that the main effect of hippocampal high-gamma was not influenced by the method of epoching, in the post-hoc analyses, we epoched the data from the end of the trial to 4.5s onward. Beamformer analyses were carried out for the high-gamma power during this time period. Then a 2 (condition: *post hidden platform* vs. *post random swimming*) × 2 (training set: 1st vs. 2nd) within-subject ANOVA analysis was performed for high-gamma power within the hippocampal and parahippocampal ROI. We reasoned that we should get similar results as those from the primary analyses, because the main time periods largely overlapped for the two epochs and the brain activities we were interested were spontaneous activities, which were not phase‐ or time-locked to any stimulus. Since the SAM beamformer can capture non-phase/time locked activities, how the data was epoched and time-aligned should not in principle influence the main results.

## Results

### High-gamma during ITI

There was a main effect of condition (F = 6.948, p < 0.05, corrected, η2 = 0.29) and a main effect of training set (F = 7.85, p < 0.05, corrected, η2 = 0.316) for high-gamma power in the right hippocampal ROI during ITI (Figure 2A). No significant effects were found for high-gamma power in the left hippocampal ROI. These results were confirmed by a whole brain 2 (conditions: *post hidden platform* vs. *post random swimming*) × 2 (training sets: first vs. second) repeated measures ANOVA analysis, which revealed a significant main effect of training set (p < 0.005, q< 0.05, FDR corrected, peak voxel in right hippocampus, Talairach coordinates: × = 18 y = -5 z = -8) and a main effect of condition (p < 0.005, q< 0.05, FDR corrected, peak voxel in right parahippocampus, Talairach coordinates: × = 27 y = -4 z = -24) (Figure 2C). No other significant effects were found in other parts of hippocampi.

**Figure 2.**
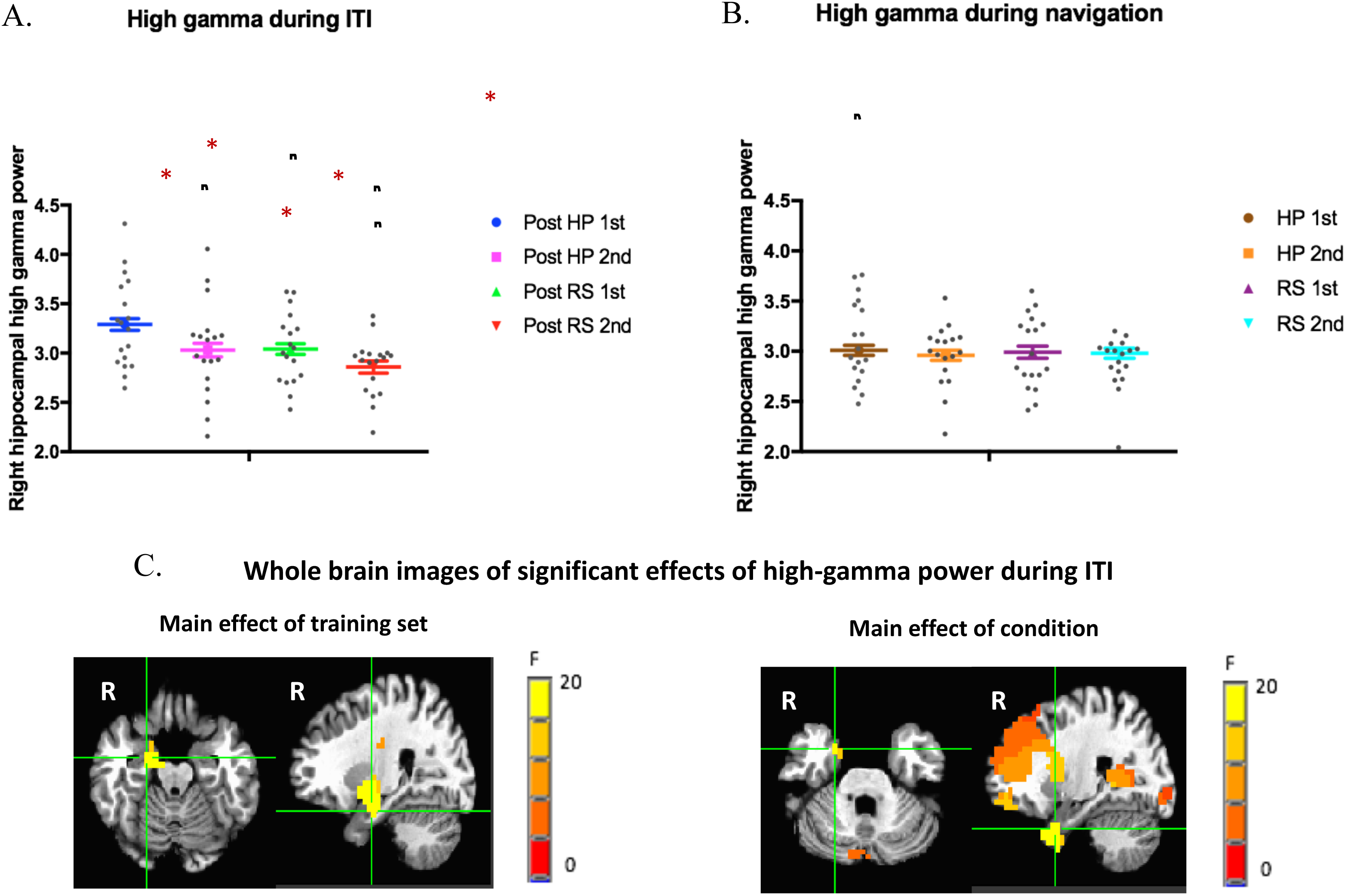
**A.** The mean of high-gamma power (pseudo-Z values) during ITI (-4.5 – 0 s) in the right hippocampal ROI regions in the first and second training set for both *post hidden platform* (Post HP) condition and *post random swimming* (Post RS) condition. **B.** The mean of high-gamma power (pseudo-Z values) during navigation period (0 – 4s) in the right hippocampal ROI in the first and second training set for both *hidden platform* (HP) and *random swimming* (RS) condition. **C.** Whole brain images of main effect of training set (peak voxel in the right hippocampus: Talairach coordinates × = 18 y = -5 z = -8) and main effect of condition (peak voxel in the parahippocampus: Talairach coordinates × = 27 y = -4 z = -24) during ITI. Error bar represents standard errors. * represents p < 0.05.

Time-frequency plots (Figure 3) show high-gamma power evolution during the ITI in the right hippocampus. The contrasts show greater high-gamma power in the right hippocampus during the ITI in the first training set (new environment) in the second training set (familiar environment); and greater high-gamma power following the *hidden platform* condition than following the *random swimming* condition. Visual inspection revealed that the greatest high-gamma increase occurred in the time window of -2.5 – 0 s in both group-averaged TFRs (Figure 3). All these results indicate that hippocampal high-gamma power exhibited the same power change pattern as hippocampal theta power during navigation in the new vs. familiar environment and high-gamma power during the ITI after navigating in the *hidden platform* condition which required more learning was stronger than that after navigating in the *random swimming* condition where learning requirement was much lower.

**Figure 3.**
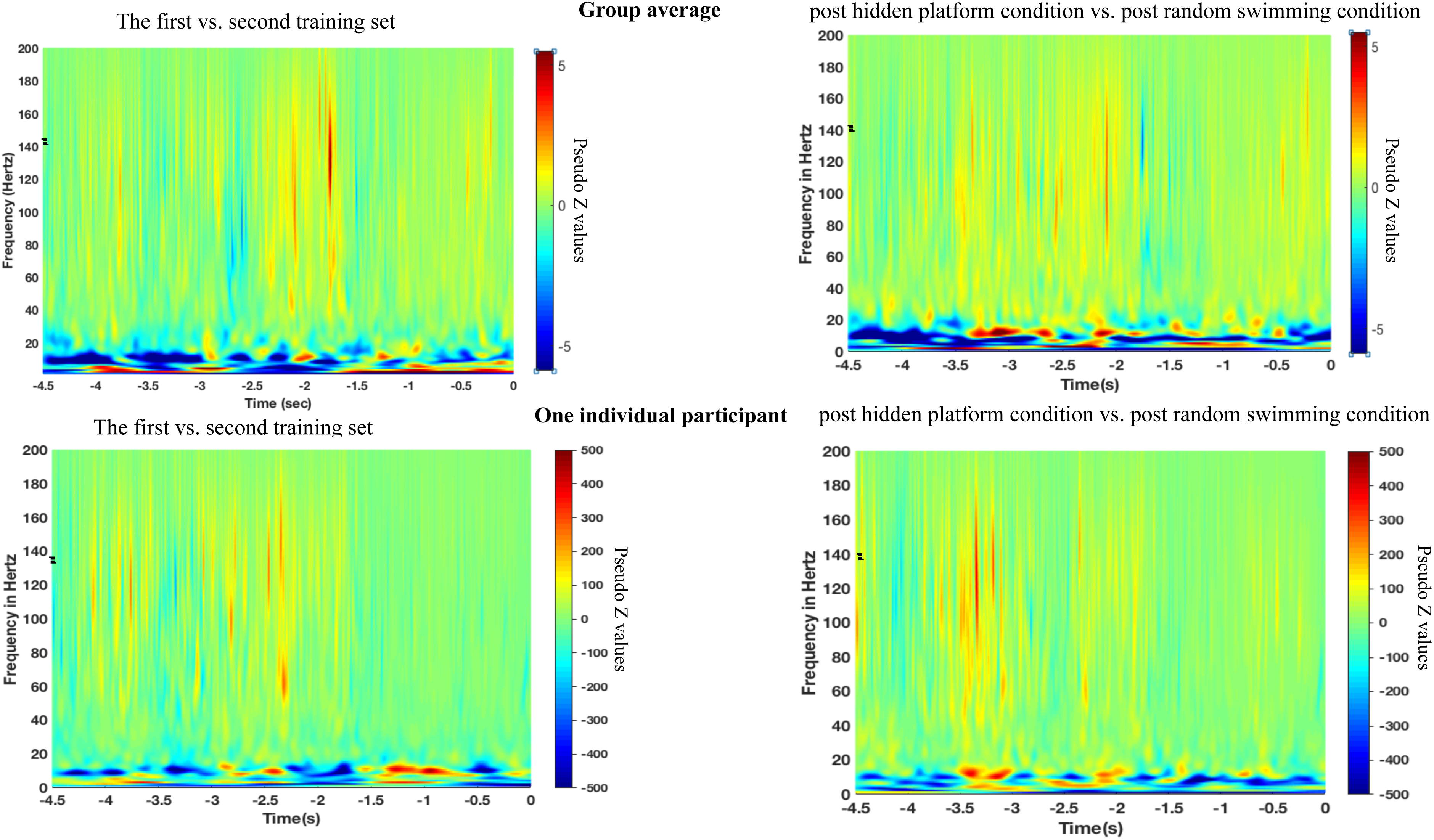
The time frequency representations (TFRs) of the peak voxel in the right hippocampus. **The upper panel** plots show the evolution of power change of right hippocampal high-gamma during the ITI in first training set relative to the second one collapsed across conditions (this shows the TFRs of the main effect of training set revealed by ANOVA analyses) and the evolution of power change of right hippocampal high-gamma during the ITI in post *hidden platform* condition relative to the *random swimming* condition collapsed across training set (this shows the TFRs of the main effect of condition revealed by ANOVA analyses). **The lower panel** shows right hippocampal high-gamma power change of one individual participant. The black rectangular shows high-gamma (80 – 140 Hz) increase during ITI as revealed by SAM beamformer analysis.

### High-gamma during navigation and its comparison with high-gamma during ITI

No significant results were found for high-gamma power during navigation (brain images not shown. no single voxel was found in the hippocampi and parahippocampi even when p = 0.05, uncorrected). Direct comparison of high-gamma power during ITI and that during navigation in the right hippocampal ROI showed that high-gamma power in the *post hidden platform* condition during ITI in the first training set was significant higher (t(17) = 3.072, p = 0.007, cohen’s d = 0.74, Figure 2A & Figure 2B) than that during navigation in the *hidden platform* condition in the first training set. No significant difference was found for the second training set (t(17) = 1.000, p = 0.329), indicating that hippocampal high-gamma increase during ITI was most apparent following navigation in the new environment.

No significant difference was found between high-gamma power in the ITI after *random swimming* condition and that during navigation in the *random swimming* condition in both training sets (t(17) = 1.66, p = 0.114 for the first training set; t(17) = -0.61, p = 0.548 for the second training set). This might suggest although after *random swimming* condition, right hippocampal high-gamma power showed the same pattern of right hippocampal theta power during navigation, the ‘replay’ effect during the ITI following navigation in a condition with low learning demands was not as salient as that following navigation in a condition with high learning demands (Eschenko & Sara, 2008; Girardeau et al., 2014). This conclusion was supported by the result of direct comparison between the high-gamma power difference between *post hidden platform* and *hidden platform* condition (Diff_H) in the first training set and the high-gamma power difference between *post random swimming* and *random swimming* condition (Diff_R) in the first training training set, which showed that the Diff_H in the first training set was significantly larger than the Diff_R in the first training set (t(17) = 2.264, p= 0.037, cohen’s d = 0.589).

## Correlation Results

### Right hippocampal high-gamma during ITI vs. navigation performance

Consistent with our hypothesis that the more the hippocampal high-gamma power increase was during the ITI after encoding the new environment, the faster the participant would learn a new location in the subsequent training set when the environment had become familiar, we found a significant correlation between the power increase of high-gamma during the ITI after *hidden platform* condition relative to that during navigation period in the *hidden platform* condition in the right hippocampal ROI in the first training set with learning rate in the second training set (r= 0.601, p= 0.004, one-tailed Figure 4A). We also found a significant correlation between the hippocampal high-gamma increase in the right hippocampal ROI with the slope of the regression line which fit the path lengths across blocks (r= -0.418, p= 0.042, one-tailed. Figure not shown). The significant correlations indicate navigators who showed higher high-gamma power during rest period after spatial learning in the new environment learned more quickly a new hidden platform location in the same environment. This is in line with results from previous literature that high-gamma during rest period after learning is associated with better subsequent memory performance (Axmacher et al., 2008).

**Figure 4.**
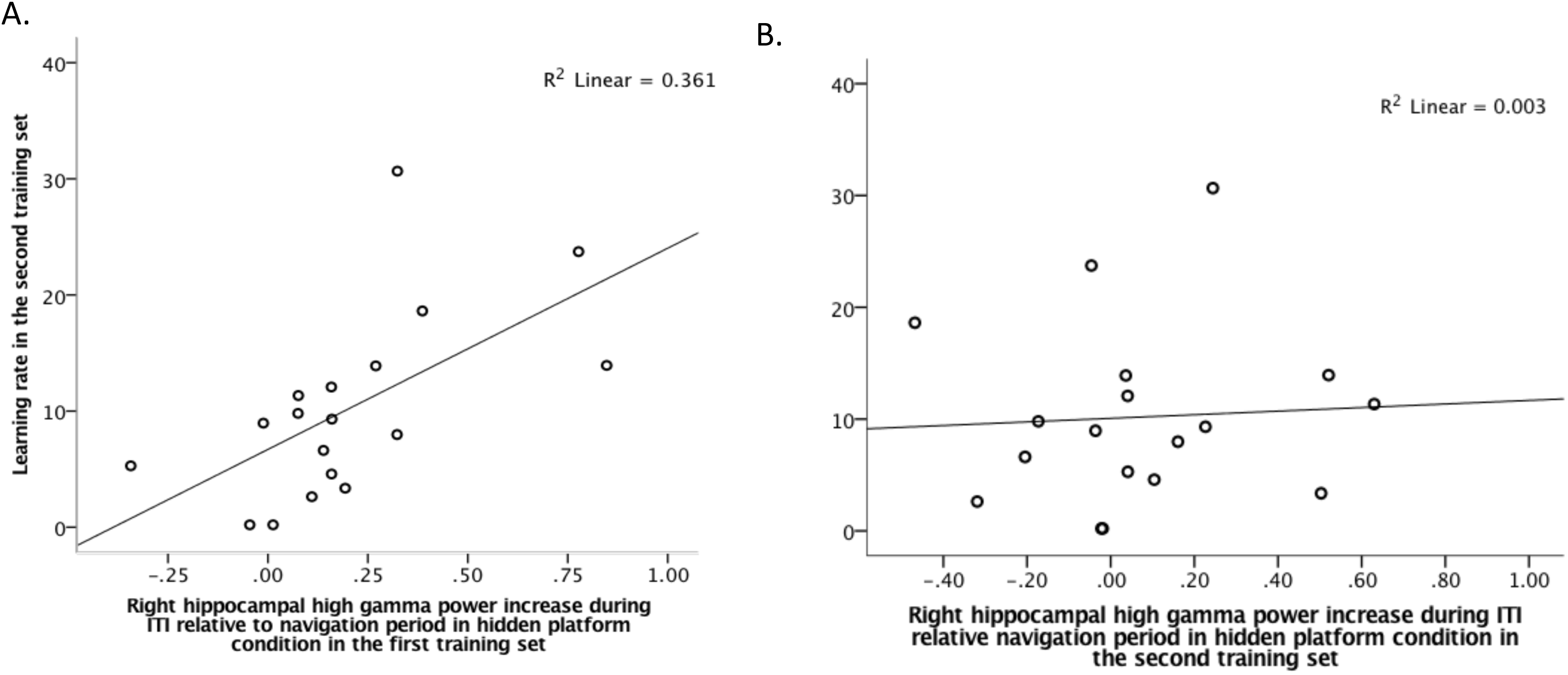
**A.** High-gamma power increase (pseudo-Z values differences) in the *post hidden platform* condition during rest relative to that in the *hidden platform* condition during navigation in the first training set (new environment) in the right hippocampal ROI (x-axis) of each participant plotted against his learning rate during navigating in the second training set (familiar environment) (y-axis). **B.** High-gamma power increase (pseudo-Z values differences) in the *post hidden platform* condition during rest relative to that in the *hidden platform* condition during navigation in the second training set (familiar environment) in the right hippocampal ROI (x-axis) of each participant plotted against his learning rate during navigation in the second training set (familiar environment) (y-axis). The y-axis of Figure 4B is the same as that of Figure 4A.

To exclude the possibility that the correlation was not learning specific and only reflected a general relationship between high-gamma power and the general cognitive processing speed across subjects, we also correlated the learning rate in the second training set in the *hidden platform* condition with the right hippocampal high-gamma power change during the ITI after the *hidden platform* condition relative to that in the hidden platform condition in the ROI in the second training set, when learning requirement decreased indexed by improved navigation performance in the second training set as shown in Pu et al. (2017). No significant correlation was found (r = 0.057, p = 0.411, one-tailed, for the correlation between hippocampal power and learning rate, r = 0.029, p = 0.455, one-tailed, for the correlation between hippocampal power and the slope of the regression line. Figure 4B). This result indicates that the correlation between high-gamma power increase after learning a new environment and subsequent learning performance in the familiar environment is functionally relevant.

### Right hippocampal high-gamma during ITI vs. right hippocampal theta during navigation

Consistent with the hypothesis that replay is proportional to encoding, we found a significant correlation (r = 0.406, p = 0.046, one tailed) between right hippocampal high-gamma power during the ITI after navigating in the *hidden platform* condition in the first training set and right hippocampal theta power during navigation. Since high-gamma power increase was most salient in the time window of -2.5 – 0s in the group-averaged TFRs shown in Figure 3, we reasoned the correlation between the right hippocampal high-gamma power in this time window with theta power during navigation would be even stronger. As expected, the correlation was significant with higher correlation coefficient (r = 0.53, p = 0.017, one-tailed, Figure 5A). Voxel wise correlation analysis confirmed that the significant effect was focal with the local maximum in the right hippocampus (Talairach coordinate of the peak voxel: × = 26, y = -17, z = -8, Figure 5C). These significant correlations support the hypothesis that ‘replay’ is proportional to encoding.

**Figure 5.**
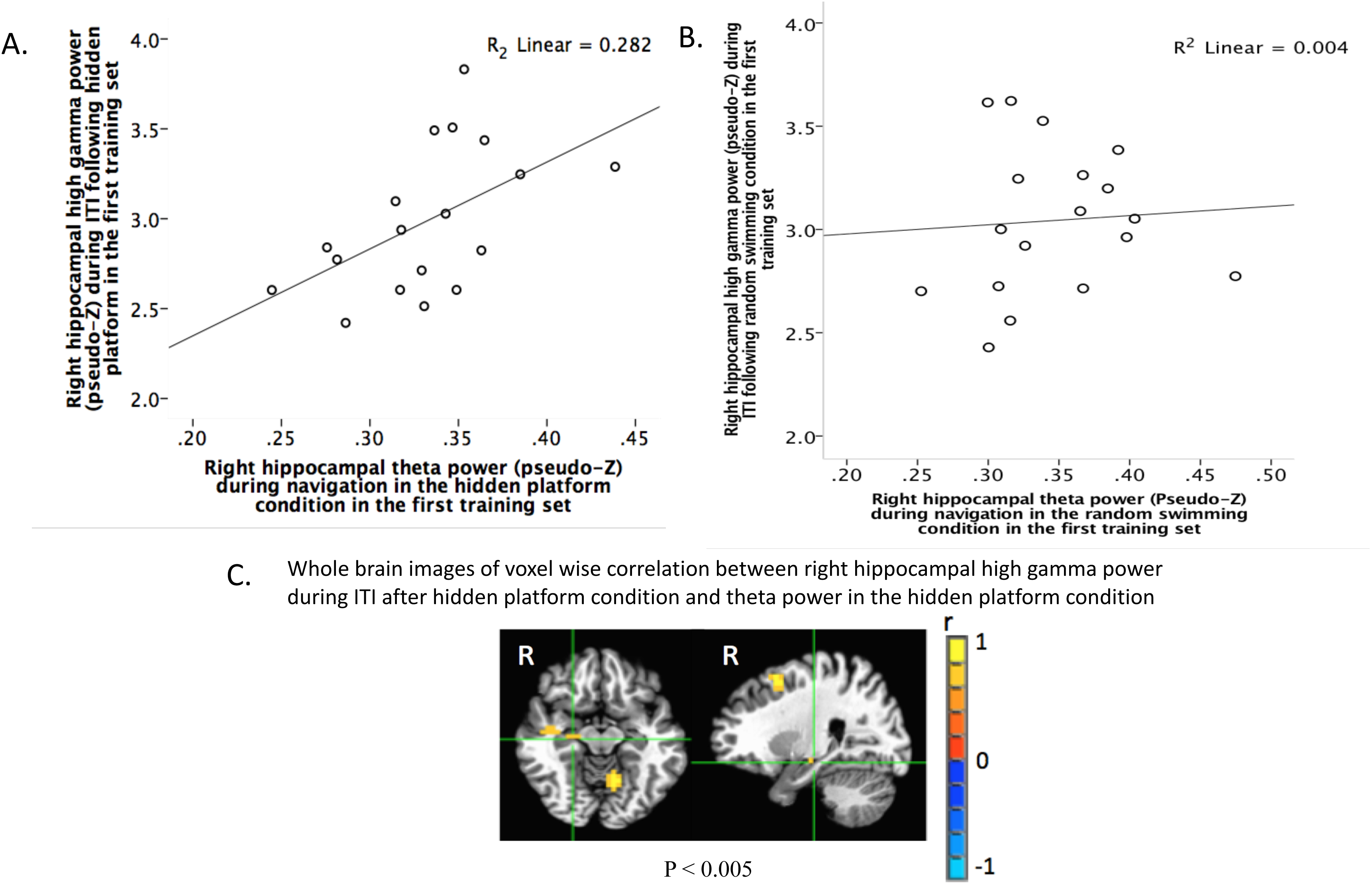
**A.** The mean of theta power (pseudo-Z values) in the right hippocampal ROI during navigation in the hidden platform condition in the time window of 1.25 – 2.25s when there was an environmental encoding effect as shown in Pu et al. (2017) plotted against the mean of high-gamma power (pseudo-Z values) in the same region during ITI of -2.5 – -0s (the time window showed the strongest high-gamma increase in the group-averaged time frequency representations in Figure 3) after navigation in the *hidden platform* condition. **B.** The mean of theta power (pseudo-Z values) in the right hippocampal ROI during navigation in the time window of 1.25 – 2.25s when there was an environmental encoding effect as shown in Pu et al. (2017) plotted against the mean of high-gamma power (pseudo-Z values) in the same region during the ITI after navigation in the *random swimming* condition. **C.** Whole brain images of the correlation between the mean of high-gamma power (pseudo-Z values) in the right hippocampal ROI in the time window of -2.5 – 0s and theta power (pseudo-Z values) in the time window of 1.25 – 2.25s in each voxel across the whole brain. Threshold is set at p<0.005 (uncorrected). The local maximum is at right hippocampus (peak voxel: Talairach coordinates × = 26, y = -17, z = -8).

No significant correlation was found between high-gamma power during the ITI after *random swimming* condition in the first training set and theta power in *random swimming* condition in the first training set (r = 0.065, p = 0.399, one-tailed, Figure 5B). Voxel wise correlation did not yield significant correlation in any voxels in the bilateral hippocampi as well. These results may indicate that the degree of faithfulness of ‘replay’ following spatial navigation in a simple environment with low encoding requirements is low, which is in agreement with findings from animal literature (Kentros et al., 2004) that faithful retrieval of a mouse’s hippocampal representation of an environment increased as task demands increased.

In the second training set where learning requirements had already decreased, no significant correlation was found between high-gamma power during ITI in the second training set and theta power during navigation in the second training set for both conditions (r = 0.078, p = 0.379, one-tailed, for correlation between high-gamma power in the *post hidden platform* condition and theta power during navigation in the second training set and theta power during navigation in the *hidden platform* condition in the second training set; r = 0.21 p = 0.201, one-tailed, for correlation between high-gamma power in the *post random swimming* condition and theta power during navigation in the *random swimming* condition in the second training set;), suggesting that the significant correlation seen above is contingent on learning.

### Post-hoc analyses

Similar effects were found for the hippocampal high-gamma power during the epoched time from the end of the navigation trial to the 4.5s forwards as for the high-gamma power epoched from -4.5 – 0s in the primary analyses above, confirming that the significant effects of high-gamma power during ITI were not influenced by the epoching method.

## Discussion

Using MEG and a virtual Morris water maze task, we investigated whether human hippocampal high-gamma power exhibited the same power change pattern as theta rhythms during navigation. We found first, that hippocampal high-gamma power after navigation mirrored the power change pattern of hippocampal theta oscillations during navigation: After navigation in the new environment, hippocampal high-gamma power was significantly stronger than that after navigation in the familiar one, and the hippocampal high-gamma power after navigating in the *hidden platform* condition where learning requirement was higher than that after navigation in the *random swimming* condition with much lower learning requirement; Second, that right hippocampal high-gamma power during ITI was correlated with right hippocampal theta power during navigation; Third that, higher right hippocampal high-gamma power after navigation in the first training set where the environment was new correlated with faster learning in the second training set where the environment became familiar.

That right hippocampal high-gamma power during the ITI after navigation exhibited the same power change pattern as right hippocampal theta power during navigation is in line with the prediction of two-stage model for memory formation (Buzsaki, 2015) and animal studies (Ambrose et al., 2016; Davidson et al., 2009; Dupret et al., 2010; Jackson et al., 2006; Jadhav et al., 2012; Karlsson & Frank, 2009; Singer & Frank, 2009, refer to Roumis & Frank, 2015 for a review) that the hippocampus is spontaneously reactivated during rest/sleep period accompanied by HFOs to promote neuronal plasticity and stabilize the newly-formed memory traces. Our finding of greater high-gamma power in the ITI after exposure to the new environment than in the ITI following the familiar environment is consistent with reports from animal studies that HFOs after navigation in a new environment are stronger, occur more frequently and are more easily detected (Foster & Wilson, 2006; O’Neill et al., 2008; Cheng & Frank, 2008; Csicsvari et al., 2007).

Our data further showed no significant high-gamma power during navigation period, when the theta rhythm was prominent (Pu et al. (2017). In addition, high-gamma power after navigation in *hidden platform* condition in the new environment was significantly greater than that during navigation in *hidden platform* condition in the new environment, which is corroborated by the idea that relative to navigation period, HFOs are more salient during the rest period when animals are disengaged from the external environment (Buzsaki, 2015). In the ITI after navigation in the familiar environment, with decreased learning requirements (indexed by improved navigation performance shown in Pu et al., 2017), no high-gamma power increase was found. These results suggest that high-gamma power increase is modulated by environmental novelty and learning requirements (Cheng & Frank, 2008). Moreover, although high-gamma power during ITI after *random swimming* trials showed a power change in the new vs. familiar environment, high-gamma increase following navigating in *hidden platform* trials during ITI relative to that during navigation in *hidden platform* trials in the new environment was significantly larger than high-gamma increase after navigating in *random swimming* trials during ITI relative to that during navigation in *random swimming* trials. This indicates that high-gamma power during rest period are stronger after navigation in a more complex environment with higher learning requirements, in agreement with the finding that replay strengths vary as a function of task demands (Eschenko & Sara, 2008; Girardeau et al., 2014).

The present results complement previous studies (Cornwell et al., 2014; Axmacher et al., 2008) by showing that the same hippocampal region used for encoding accompanied by low frequency oscillation exhibited a similar power change pattern accompanied by high-gamma oscillations during ITI after encoding, thus supporting the role of hippocampal high-gamma in replay of newly learned experience. These results also support previous fMRI studies (Deuker et al., 2013; Gruber et al., 2016; Peigneux et al., 2006; Staresina et al., 2013; Tambini & Davachi, 2013; Tambini et al., 2010; Vincent et al., 2006), which showed experience-dependent reactivation during wakefulness or sleep and provides a neurophysiological mechanism underlying the reactivation. The current results may also help explain why some fMRI studies have not found hippocampal reactivation during rest/sleep period after learning (e.g., Deuker et al., 2013, they found visual area showing a “replay” effect instead of the hippocampus). Because as shown in our data, hippocampal reactivation occurs immediately after each learning trial and decays over learning, during which information is being transferred to neocortex (Colgin, 2016; Kudrimoti et al., 1999). Therefore, hippocampal reactivation might not be as strong and salient during rest/sleep after a long period of learning as that immediately after each learning trials on a faster timing scale. Further study could investigate how persistent high-gamma-related hippocampal replay is.

HFOs measured using MEG are easily confounded by muscle artifacts. However, source localization algorithms based on spatial filters can differentiate cognitive processing source from cortical or subcortical areas with artifactual sources (Dalal et al., 2011). Several checks support our contention the high-gamma effects observed in the current study are not artefactual. First, the analyses were performed during the inter-trial period when participants were instructed to rest quietly and to minimize movement. Second, there was no significant high-gamma effect during active navigation epochs which are more likely to be contaminated by muscle artifacts because participants pressed buttons to move in the virtual pool. Third, the spatial map of the effect was focal and localized unilaterally to right hippocampus, and TFRs showed relatively narrow band power changes. Muscle artifact, in contrast, tends to span large spatial regions and broad frequency ranges (from 30 to 200 Hz or higher; Muthukumaraswamy, 2013).

Consistent with our hypothesis that replay is proportional to encoding, our data showed that high-gamma power during the ITI after navigating in the *hidden platform* condition in the new environment was positively associated with theta power during navigation in *hidden platform* condition in the new environment, when learning requirement was strongest. Together with the finding that the same hippocampal region used for encoding was reactivated during rest period, the correlation provides further evidence that high-gamma during rest is modulated by previous learning experience to accurately reinforce the newly-formed labile memory traces (Davidson et al., 2009; Skaggs & McNaughton, 1996; Wilson & McNaughton, 1994). A comparable correlation was not found for familiar environments with decreased learning requirements, indicating that the correlation seen in the new environment is learning induced, rather than an intrinsic relationship between gamma and theta power. The random swimming condition, where learning requirements were low, showed no significant correlation between theta and high-gamma in both training sets, suggesting although replay is automatic, the degree of replay faithfulness may vary as a function of encoding requirement. This is corroborated by the findings of a rodent study (Kentros et al., 2004), showing that the faithful retrieval of a mouse’s hippocampal representation of an environment increases as task demands increase and place cell stability tightly covaries with attention to the available spatial cues.

Further, we observed that right hippocampal high-gamma power increase following *hidden platform* condition in the first training set relative to that during navigation in this condition in the same training set correlated with learning rate in the *hidden platform* condition in the second training set. This correlation provides strong evidence for the argument that the functional role of right hippocampal high-gamma reactivation is memory consolidation. Navigators with higher right hippocampal high-gamma power during the ITI after navigation in a new environment learned more quickly a new location of the hidden platform in the second training set where the environment was the same and become familiar, indicating right hippocampal high-gamma power is related to consolidation of the newly learnt environment. Good consolidation of the environment to form a cognitive map of the surroundings can facilitate flexible navigation to any place in the same environment (Wolbers & Hegarty, 2010). This correlation is also consistent with observations from human fMRI studies showing that reactivation strength of the hippocampus predicts subsequent memory performance (Bergmann et al., 2012; Peigneux et al., 2006). The direction of the correlation is also consistent with the results of Axmacher et al. (2008) and Cornwell et al. (2014), documenting that stronger high-gamma power corresponded to better memory performance. No correlation was found between high-gamma power in the second training set (where consolidation requirement decreased) with learning rate in this training set, indicating the significant correlation seen above is learning specific.

It is noteworthy that the effects described in the current study were confined to right hippocampus. In our previous findings, navigation-related theta effects were found in both left hippocampus (implicated in binding the external cues to the platform location) and right hippocampus (associated with encoding the environment to form the cognitive map of the space). While non-significant results do not confirm the null hypothesis, this is consistent with the notion that replay is selective, such that not every aspect of learning would be replayed (Deuker et al., 2013). However, the mechanism underlying this replay selection is still unclear. Reactivation during awake rest and sleep is more complicated than expected (Buzsaki, 2015). O’Neill et al. (2006) has shown the replay might not necessarily be location dependent. Recent studies (Gupta et al., 2010; Wu & Foster, 2014) have pointed out the view that the function of replay is helping construct a Tolmanian cognitive map of the environment, which would result in flexible routes to the goal location on subsequent trials. This might help explain why only the right hippocampus region, which is associated with environmental learning was reactivated in our data. Also, the present work provide new insights into the reason why right hippocampus is believed to be important in spatial cognition in general (Cornwell et al., 2010; Jacobs et al., 2009; Maguire et al., 1998; Nedelska et al., 2012). More research is needed in the future to investigate the selective nature of hippocampal replay.

In sum, using a highly translational experimental task and non-invasive MEG measurements, we show during the inter-trial period immediately following spatial learning, human hippocampal high-gamma activity is evident and plays an important role in replay of the newly-learned information. These findings advance our understanding of the neurophysiological mechanisms and timing of hippocampal reactivation after learning for memory consolidation in humans.

## Acknowledgements

This work was supported by Australian Research Council Grants CE110001021, DP1096160. The first author was financially supported by the China Scholarship Council and Macquarie University. We thank Dr. Suresh Muthukumaraswamy for helpful discussions about muscle artifacts and gamma activity. The authors declare no competing financial interests.

